# Twist and Snout: Head and Body Morphologies Determine Feeding Kinematics in Substrate-Biting Fishes

**DOI:** 10.1101/2025.05.28.656629

**Authors:** Tal Perevolotsky, Jacob M. Brotman-Krass, Yarden Ratner, Yael Avigad, Adam P. Summers, Cassandra M. Donatelli, Roi Holzman

## Abstract

Across teleosts, feeding by biting substrate-attached prey has evolved multiple times and is associated with convergent morphologies that include a deep body and an elongated, tapered head. However, the functional role of these morphologies in substrate-biting fish is not established. Here, we tested the hypothesis that these morphologies function as control surfaces that affect feeding kinematics during biting. To test this hypothesis, we used simplified physical models of substrate-biting fish and examined the role of head, body, and fin morphology in determining feeding kinematics that facilitate the removal of substrate-attached prey. Models simulated the swift lateral movement of the head, previously documented in species biting substrate-attached algae. Using models that capture the natural morphological variation of biters, we tested (i) how different head morphologies affect the speed of the head and (ii) how different body morphologies affect the stability of the body during head movements. We found that the moment of inertia (MOI) of the head and body explained most of the variation in head speed and body displacement. A decrease in head MOI resulted in faster lateral head movements, known to facilitate removal of attached prey. An increase in body MOI, relative to that of the head, stabilized the lateral displacement of the body during bites. Overall, our results suggest that the laterally compressed bodies and tapered snouts function as control surfaces during feeding in substrate-biting fish. We propose that a selective pressure to extend the lateral surface area underlies the prevailing morphological convergence of biting reef fishes.

## Introduction

Fish display an astonishing trophic diversity, feeding on practically any organic food source found in their aquatic surroundings [Hiatt and Strasburg, 1960, Parravicini et al., 2020, Ng et al., 2024]. This trophic domination is attained through numerous morphological adaptations, supporting various feeding mechanisms that allow fish to procure a diversity of prey types [Wainwright et al., 2002a]. One of the most common mechanisms supporting this diversity is the ability of fish to obtain prey by biting. Biting is defined as a feeding behavior that involves the direct contact and manipulation of prey by the fish’s jaws and teeth [Wainwright et al., 2002a, Corn et al., 2021]. Biting is usually required when feeding on large, tough, or armored prey, or when the prey is attached to a substrate. It is a prevalent feeding behavior that encompasses species from a diversity of trophic levels, morphologies, habitats, and phylogenetic origins [McGee et al., 2016, Borstein et al., 2019, Floeter et al., 2018, Corn et al., 2022]. Biting is especially common on coral reefs where, since the early Cenozoic, reef-associated biting lineages have rapidly evolved, leading to over a ten-fold increase in their overall frequency. As a result, biters comprise 40% of modern reef species and some of the most prominent reef-associated families are predominantly biters, including Labridae, Scaridae, Acanthuridae, Siganidae, Chaetodontidae, and Pomacanthidae. [Corn et al., 2022].

Biting fish display an immense diversity in diets and feeding behaviors, with prey varying dramatically in size, ecology, and mechanical properties. For example, invertivore behavior ranges from picking small crustaceans from the sand to crushing the exoskeleton of armored Echinoderms. Herbivores feed on algae ranging from fine, millimeter-long turf to large, leathery macroalgae. Some Corallivores display high dexterity in picking individual coral polyps while others excavate deep into their calcium-carbonate skeleton [Tebbett et al., 2017, Bellwood and Choat, 1990, Turingan, 1994, Streit et al., 2015, Cole et al., 2008].

Biters have been the focus of many studies aimed at characterizing their diverse ecological niches, which often hinge heavily on dietary preferences. Keystone herbivorous parrotfish, for example, are usually classified into ecologically relevant ‘functional groups’, namely scrapers, excavators, and browsers. This classification is based on the degree of their interaction with the substrate and gut content, but also extended to the morphology of their jaws, teeth, and jaw muscles [Bellwood and Choat, 1990, Green et al., 2009, Bonaldo et al., 2014, Nanami, 2016, Tada et al., 2017]. Studies aimed at characterizing the feeding kinematics of biters have focused mainly on large carnivorous biters, while the feeding kinematics of reef herbivores have received much less attention [Lemberg et al., 2019, Porter and Motta, 2004, Grubich et al., 2008, Ferguson et al., 2015]. Specifically, the link between biting kinematics and the ecological effect of herbivory is poorly understood [but see Mihalitsis and Wainwright, 2024, Perevolotsky et al., 2020].

From a biomechanical perspective, the major body of work concerning biting biomechanics has focused mainly on (1) biting force - mostly through models of force transmission [Westneat, 2003, 2004, 1994] (2) functional morphology of the teeth and the point of contact with the prey [Galloway et al., 2016, Purcell and Bellwood, 1993, Nanami, 2016, Grubich et al., 2008, Ferguson et al., 2015, Cohen et al., 2023, Whitenack and Motta, 2010] (3) and innovations and novel morphologies of the above [Konow and Bellwood, 2005, 2011, Gibb et al., 2015, Price et al., 2010, Olivier et al., 2021, Evans et al., 2023]. The kinematics of biting have primarily been studied in comparison to suction feeding. As a result, these studies focus on gape and cranium morphology and kinematics [Martinez et al., 2022, Corn et al., 2021]. It is less clear which functional traits determine the ability of biting fish to procure different prey types and how morphology is linked to biting performance beyond bite force.

In-situ high-speed recordings of Acanthurid fish feeding in the natural reef environment revealed that the process of removing substrate-attached prey is much more intricate than previously thought, including the synchronized kinematics of multiple body parts, along with the gape cycle [Perevolotsky et al., 2020]. This work focused on feeding kinematics of two species of *Zebrasoma*, a prominent reef herbivore, grazing on short filamentous algae in the shallow reef. *Zebrasoma* are characterized by a deep laterally compressed body, protruding snout, and a small mouth equipped with multi-cusped teeth [Randall, 1955]. Perevolotsky et al. [2020] measured fish whole-body kinematics, the force they exerted on the substrate, and the amount of algae removed during feeding. *Zebrasoma* displayed highly stereotypical feeding kinematics that included closing their mouth and grabbing the algae, followed by a fast, forward swing of the pectoral fins and a lateral movement of the head. Hereafter, we refer to this biting mechanism, in which the prey is procured by exerting a pulling force using the entire body, as ‘pulling from the substrate’. In this feeding mode, bites characterized by faster body kinematics removed more algae from the substrate. In other words, the speed at which fish backed away from the substrate (influenced by pectoral fin and head flick speed) determined the pull force they exerted, which in turn determined the amount of algae removed.

Interestingly, the sideways movement of the head (‘head-flick’), occurring at the end of the bite, was previously documented in laboratory experiments [Purcell and Bellwood, 1993, Ferry-Graham et al., 2002, Konow and Bellwood, 2011, Mihalitsis and Wainwright, 2024] and in observations of natural feeding behaviors [Jones, 1968, Clements and Bellwood, 1988, Motta, 1988, Streit et al., 2015, ;Figure 1] in a diversity of species from families that include Acanthuridae, Labridae, Chaetodontidae, Kyphosidae, and Pomacanthidae (Figure 1). The prevalence of head-flicking behavior across biting species suggests it is an integral part of the feeding repertoire of fish feeding from the substrate. Furthermore, it is functionally important to feeding performance as shown in [Perevolotsky et al., 2020].

**Fig. 1.**
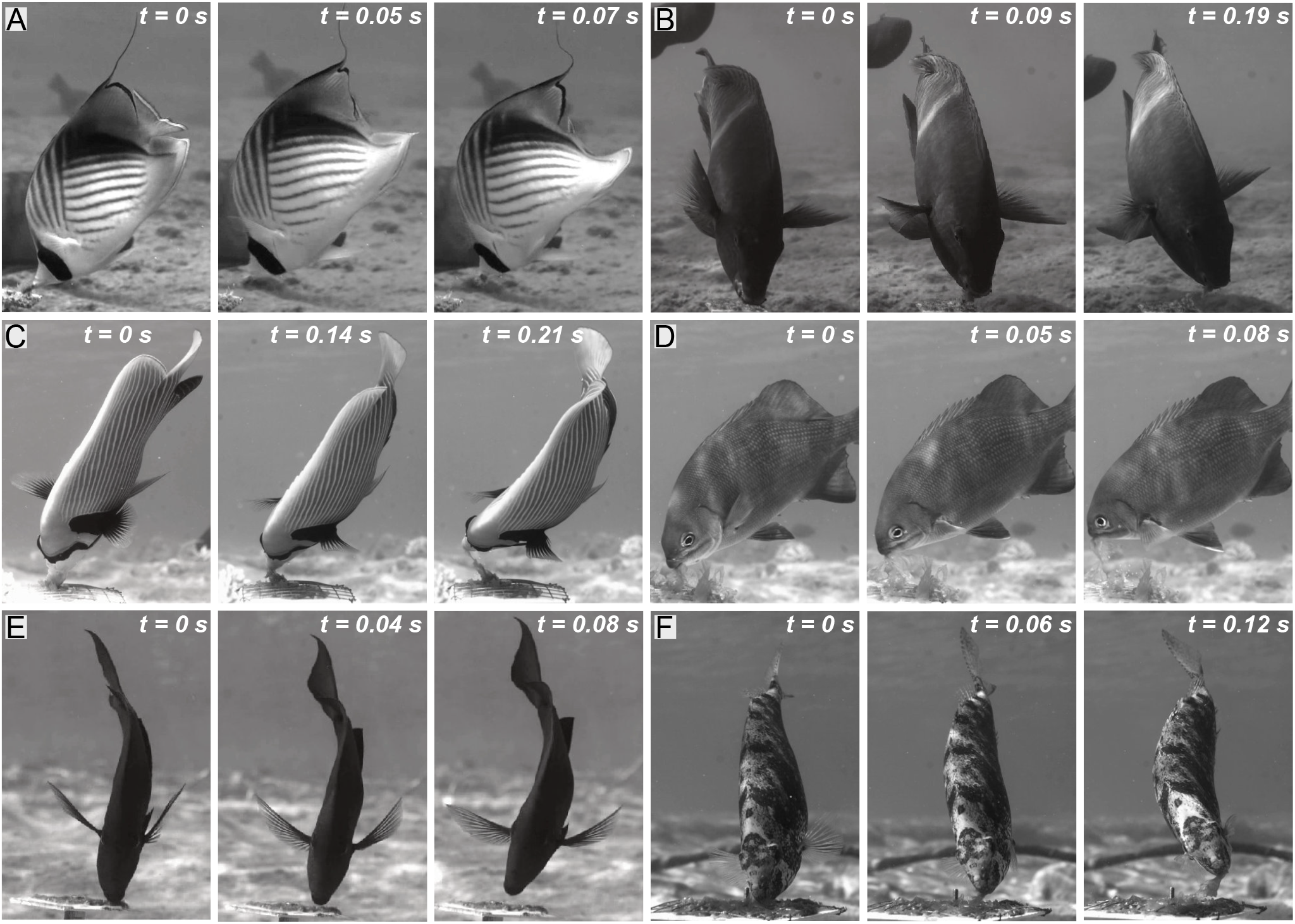
Head flicks are commonly used by substrate-biting fish. Videos of in-situ feeding events [using the system described in Perevolotsky et al., 2020] revealed that species from different families and trophic niches all perform a fast lateral movement of the head at the end of the bite, augmenting the pull force used to remove attached prey. Species and their respective prey include A) *Chaetodon auriga*, live anemone, B) *Scarus fuscopurpureus*, algae and fish meal gel, C) *Pomacanthus imperator, Ulva lactuca*, D) *Kyphosus vaigiensis, Ulva lactuca*, E) *Acanthurus nigrofuscus*, filamentous algal matrix, F) *Siganus rivulatus, Ulva lactuca*. For each photo sequence, the timing of events (in seconds) includes: 1) The beginning of the flick, as the mouth closes on the prey (t = 0), 2) mid-flick, before the prey detaches completely, and 3) flick end, when the head reaches maximum lateral displacement. Flick duration ranged from 0.07s in *Chaetodon auriga* to 0.21s in *Pomacanthus imperator*. Each face of the feeding plate in A,B, and E is 5cm and 10cm in C, D, and F.

The body of species observed biting from the substrate *in-situ*, in the natural coral reef, remained remarkably stable during feeding bouts, despite the vigorous lateral motion of the head [personal observations and data from Perevolotsky et al., 2020, ;Figure 1]. Body stability should be important in enabling the fish to perform multiple bites from the same confined area, without redirecting and swimming back to the same spot. We therefore suggest that fish that pull from the substrate maximize two performances: (1) augmenting the speed of the mouth with respect to the prey, increasing prey removal per bite while (2) minimizing the movement of the body with respect to the prey, increasing stability and allowing for multiple bites per feeding bout. Moreover, we suggest that the head and body morphologies of pullers should affect their kinematics during head flicks.

We use fundamental mechanical principles to predict the links between head flick kinematics and the morphology of the fish. Specifically, Newton’s second law (applied for rotational movement) predicts that the angular acceleration of the head about its axis will depend on the torque (applied by the body muscles) and the moment of inertia (MOI) of the rotating head. MOI, in turn, sums up the distribution of mass weighted by the square of its distance from the axis of rotation (Equation (1), in methods). In other words, the MOI can potentially be used as a morphological index that predicts head flick kinematics.

Furthermore, the conservation of angular momentum dictates that the angular momentum of the head and that of the body should be equal and opposite, given no external torques. Hence, the rapid movement of the head should also drive rotation and translation of the fish’s body [Lighthill, 1970]. The degree of body rotation depends on its resistance to rotation; specifically, the ratio between body MOI and head MOI determines how fast the body must rotate to compensate for a given head rotation. We hypothesize that balancing the lateral movements of the head is achieved through expansion of the dorsal, anal, and caudal fins, along with the body surface itself. These lateral morphologies [aka “lateral profile” Webb, 1978, 2005, Fish and Lauder, 2017] should function as stabilizing control surfaces that resist rotation.

Accordingly, we hypothesize that the two (possibly contrasting) functional demands imposed on head-flicking pullers, i.e., increasing head angular rotation while decreasing body displacement, are facilitated by two morphologies: 1) Head morphologies that are characterized by low moment of inertia will facilitate faster lateral movements of the head (possibly augmenting pull force); and 2) body morphologies that are characterized by a high moment of inertia relative to the head, will reduce body displacement, facilitating stability during feeding bouts. We test these hypotheses using an experimental system simulating head flicks in simplified mechanical fish models. Varying the model’s head and body morphologies allowed us to examine links between morphology and kinematics under controlled conditions.

## Methods

### Fish species

We constructed physical models based on seventeen substrate-biting coral reef species, belonging to seven iconic reef radiations [Corn et al., 2022]. Species were chosen to represent a diversity of body, fin, and head morphologies; as well as a range of trophic niches, including herbivores, corallivores, and general invertebrate feeders (Supplementary Table 1). Species included *Chaetodon auriga* and *Chaetodon melapterus* (Chaetodontidae), *Zebrasoma xanthurum, Acanthurus fowleri, Acanthurus nigrofuscus*, and *Naso unicornis* (Acanthuridae), *Siganus vulpinus* and *Siganus rivulatus* (Siganidae), *Anampses caeruleopunctatus, Iniistius pavo*, and *Calotomus viridescens* (Labridae and Scaridae), *Stegastes lacrymatus*, and *Pomacentrus aquilus* (Pomacentridae), *Centropyge multispinis*, and *Pomacanthus imperator* (Pomacanthidae), *Kyphosus vaigiensis*, and *Microcanthus strigatus* (Kyphosidae). All seventeen species were observed performing ‘head flicks’ during feeding either in-situ on the reef, or in videos of feeding behaviors in aquaria (Figure 1, Supplementary Table 1). We focused on fish that forage by active swimming. Our sample did not include other common substrate-biters such as Gobbies and Blennies that contact the substrate with their body and fins during feeding.

### Physical models

In order to isolate and compare the effect of head shape alone, all models were constructed to have an identical projected head area. This area was 30 cm^2^, corresponding to the projected head area measured from an adult *Zebrasoma xanthurum* with a total length (TL) of 20 cm. We note that a total length (TL) of 20 cm is common for this species [Randall, 1955, Lieske and Myers, 2004]. *Zebrasoma* was chosen as a reference because it represented the average TL of our sampled species. In order to create models with the same head area, photos of adult specimens downloaded from Fishbase [Froese and Pauly, 2024] were resized in a photo editing software (Affinity Designer) until their head area reached the desired 30 cm^2^. As a result, some species’ morphologies had to be enlarged (compared to their common size) to reach the desired head area, while others had to be reduced. Since the resizing process was performed on the entire photo, the original ratios between the head area and the combined area of the body and fins were maintained. This resizing resulted in models with equal projected head areas and different head shapes, along with lateral profiles (body and median fins) that differ in both shape (following the natural variation) and projected areas (due to resizing). It is important to note that our resizing process consisted a linear geometric transformation, meant to enable an informative comparison between shapes. It was not intended to reproduce an ontogenetic growth process. Hence, we do not presume that larger models represent a later ontogenetic stage or that smaller models represent an earlier one. Similarly, we did not try to capture the natural morphological variation within each species, which in some species can change depending on ontogeny and sex. In general, our inferences do not rely on how well the physical model represents the size and growth of each species, but are based on a comparative framework isolating the effect of shape differences between models.

Fish outlines were traced from the resized photos and included the head, body, median, and pelvic fins (Figure 2). We focused on the effect of the median control surfaces [Webb, 1978] and did not include the pectoral fins, for which we could not replicate the intricate kinematics during feeding strikes.

**Fig. 2.**
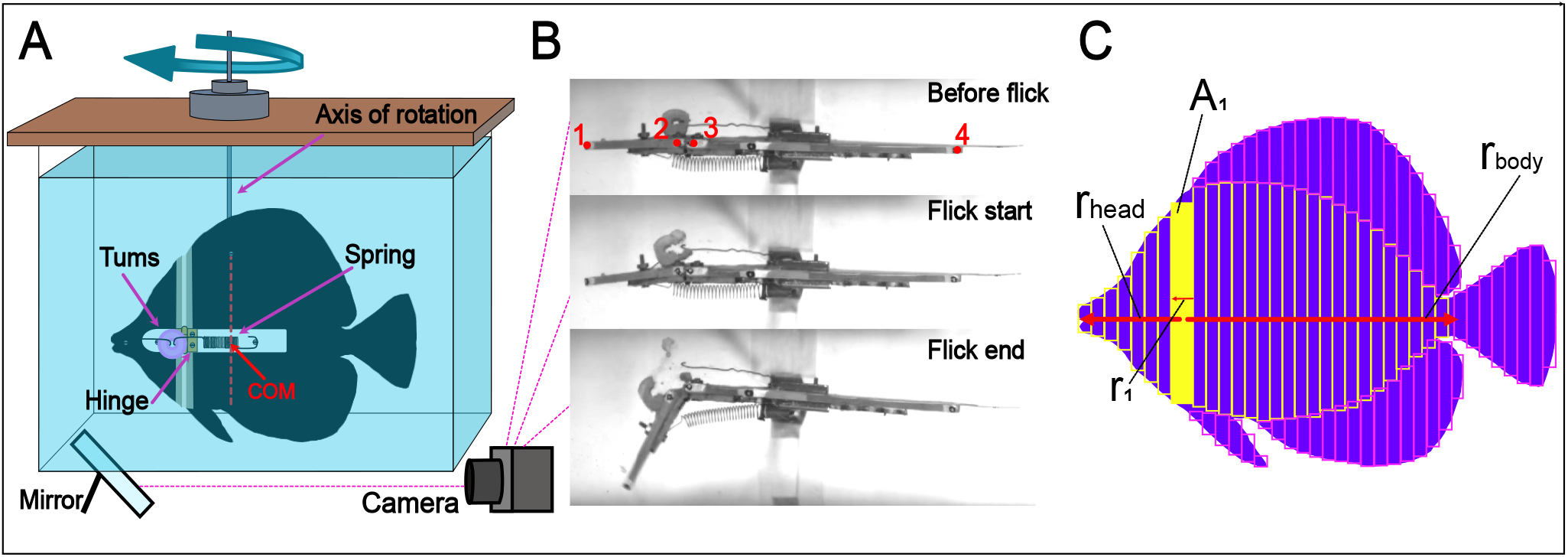
Experimental setup. A) models performed simulated head flicks in an aquarium, mounted on a rod which allowed for unconstrained rotation around their center of mass (COM). Setup illustration is not to scale. B) models were filmed from below for kinematic tracking in videos capturing their position before the latch dissolved and ending when the head reached and angle of 45^°^. Total length of the *Chaetodon auriga* model pictured is 27cm. C) The moment of inertia was calculated by measuring the surface area (e.g. *A*_1_) and the distance from the axis of rotation (e.g. *r*_1_) of segments along each morphology (here, yellow segments represent body and head morphologies and pink represent fin morphologies). Measurements were then incorporated in the MOI equation, along with the relevant material mass (see eq. (1), in methods)

Focusing on the effect of lateral morphologies, the width of the body and head was minimized and shared by all models, regardless of the natural width of their respective species. The head and body of the models were separately cut out using a CNC machine from 4 mm foamed PVC sheets. Sheets of this width were strong enough to resist flexion while moving in our experimental setup, simulating the rigidity of fish bodies while minimizing the model’s width. The dorsal, pelvic, anal, and caudal fins were made from a 0.5 mm flexible PVC sheet. Fin sheets were flexible enough to bend and flex during the model’s movement in water, while strong enough to maintain their shape and not warp. Though the ratio between body width and fin width in our models does not represent the natural ratio, the mechanical properties of model sections are (by-and large) representative of the material properties of each morphology.

Body and fin sheets were glued together using cyanoacrylate glue. The head and body were connected by a metal hinge that allowed the head to rotate freely to the left, defining the head’s axis of rotation. Loose tape was used to cover the gap between the body and head, creating a flexible joint that prevented the passage of water during flicks without hindering head rotation. Using a designated plastic frame, which was transferable between models, a metal spring was attached between the middle of the head and the center of mass of the model (Figure 2A). A latching mechanism was constructed using a dissolvable calcium bicarbonate tablet (TUMS), which was sanded to form a toroid with a square cross-section of 3 by 3 mm. The tablet was hooked between the head and body of the model using thin metal hooks located on the plastic frame, holding the spring in tension and the body and head lined up at an angle of 0^°^. When inserted to the water, the sanded tablet would dissolve and fail, abruptly releasing the spring to pull the head sideways, generating rapid head rotation (Figure 2B), that simulated the head flick. Dissolving time of the TUMS averaged ∼ 90 seconds. The movement of each model was restricted to rotation about the vertical axis (yaw) through the model’s center of mass. This was done by mounting the model on a vertical metal rod passing through the model’s center of mass, and connected via a metal bearing to a frame above the aquarium. For each model, we acquired 6-10 head flick videos. Kinematic variables from these trials were averaged for each model for a final sample size of N=17 morphologies (species). The angular speed of the models ranged 600 − 1150^°^*s*^−1^, about 2-3 times faster than natural *Zebrosoma* head flick speed [Perevolotsky et al., 2020].

### Video analysis

Models were placed in a 60×30×40 cm aquarium and filmed using a high-speed camera (Photron Fastcam SA6) at 250 frames per second. A mirror was placed under the tank at an angle of 45^°^, providing a ventral view of the model (Figure 2A). The camera was triggered manually upon latch release. Saved sequences started ∼10 frames prior to the time of latch release, and spanned the time it took the model to come to a complete stop. Time *t*_0_ was defined as the onset of head movement.

Four landmarks were physically marked on the models and digitized for each video: 1) the anterior tip of the head, 2) the posterior tip of the head (adjacent to the head-body joint), 3) the anterior ventral tip of the body (on the opposite side of the head-body joint), and 4) the posterior tip of the body (Figure 2B). From these landmarks we calculated the angle between the head and body at each frame. Using landmarks 3 and 4, we calculated the earthbound angle of the body in each frame. For each head flick video, we calculated (1) the angular speed of the head (^°^ s^−1^), based on the time (s) it took the model to reach an angle of 45^°^ (hereafter *t*_*end*_) and (2) the angular displacement of the body (^°^), calculated as the difference between the angle of the body at *t*_0_ and *t*_*end*_.

### Model’s morphology and MOI calculations

The moment of inertia (MOI) of a given morphology depends on the distribution of its mass along the axis of rotation. We calculated the MOI for the head and body by integrating the MOIs of segments along their longitudinal axes (Figure 2C), using the following equation:

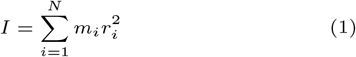

Where *I* is the moment of inertia of the morphology of interest (i.e. head, body and the fins), *r* is the distance of the segment’s centroid from the rotation axis, and *m* is the mass of the segment. We calculated MOIs from lateral photos of the models. We used ImageJ [Schindelin et al., 2012] to mark rectangles on the head, body, and fins of each model. Rectangles were oriented with their long axis parallel to the axis of rotation, each rectangle spanning the height of the morphology at the measurement point (Figure 2C). Rectangle width was set to ∼ 5 mm for the head and fins and ∼ 7 mm for the body. The number of rectangles varied according to the model’s aspect ratio. Rectangle mass was calculated as the product of its projected area * PVC weight per cm^2^. Material weight was 1.44 g*/*cm^2^ for the head and body and 0.475 g*/*cm^2^ for the fins.

### Analysis

The angular speed of the head was used to test our first hypothesis, i.e. that models with lower head MOIs would be characterized by faster flicks. This prediction is based on the relationship between torque and MOI, described by the following equation:

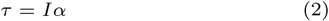

In our models, the torque *τ*, which is applied by the spring on the head, is constant; hence, a decrease in the MOI of the head will increase its acceleration, *α*. Here we chose to measure the angular speed of the head, rather than its acceleration. We argue that the speed of the head flick is correlated with its angular accelerations, because the motion starts from zero speed and develops rapidly over a finite and limited excursion (*θ*_*head*_ = 45^°^). In other words, obtaining higher flick speeds over a similar excursion requires faster acceleration. Furthermore, we elected to report flick speeds, rather than acceleration, because this metric is commonly measured in studies of fish kinematics (and other animals) and was related to ecological performance [Perevolotsky et al., 2020].

Our second hypothesis was that the morphologies of the lateral profile (body surface, dorsal, anal, and caudal fins) affect the model’s stability during flicks. The conservation of angular momentum dictates that

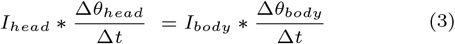

It follows that, given a constant change in head angle (*θ*_*head*_ = 45^°^) and a known flick time Δ*t*, the change in body angle *θ*_*body*_ during the flick will depend on the ratio between the head and body MOI. In other words, morphologies with low head MOI and high body MOI will maximize body stability (minimize body displacement) during head flicks.

To test the first hypothesis, we regressed the angular speed of the head (the dependent variable; averaged for each species across all trials) with the moment of inertia of the head (the independent variable).

Similarly, to test the second hypothesis, we regressed the angular displacement of the model during the flick (dependent variable; averaged for each species) with the ratio between head MOI and the combined MOI of the body and median fins, i.e., lateral MOI (this ratio is hereafter referred to as relative MOI). Statistical analysis was performed using R software version 4.4.3 [R Core Team, 2024].

### Statistical approach

Since the models used in our experiment are based on real species, some degree of morphological similarity between species can be attributed to their shared history. We therefore account for shared history when examining the correlations between morphologies of different control surfaces, in different species. This is because the evolutionary-driven patterns are the main interest of this analysis. Accordingly, we used phylogenetically corrected Pearson correlation test, using the function *cor phylo*, implemented in the R package *phyr* to test the correlation between control surfaces. In this test, the phylogenetic signal for each variable is estimated from the data assuming that variate evolution is driven by a Ornstein-Uhlenbeck process. Thus, the function allows the estimation of a phylogenetic signal in multiple variables while incorporating correlations among variables. We used a published time-calibrated phylogeny [Rabosky et al., 2018], which we pruned to the species in the study.

However, when examining the correspondence between morphology and kinematics, we chose against using a phylogenetically informed analysis. This is because the models are powered artificially, using the same mechanism that is agnostic to the evolution of morphology, muscle physiology, and its control. Hence, the species’ identity does not explain the variation in the “physiology/behavior” of the model i.e. the forces that move the body and the animal’s control over them. We feel that a (simple) linear regression is most adequate to isolate the relationship between morphology and kinematics, which is the result of the spring driving force and the physical forces that resist it (which stem from the morphology). Critically, there are no unmeasured variables that could be accounted for by the shared history of the species.

## Results

### Control surfaces evolve in a correlated manner

The MOI of the head ranged ∼2 fold, from 1969-3534 g cm^2^, despite having the same projected area. That is, this variance resulted only from changes to the shape i.e. the distribution of weight along the model’s head. The MOI of the body spanned ∼4 fold, from 124000-468000 g cm^2^. This variance resulted from differences in both shape and size.

Across our models, the control surface that contributed the most to the total lateral MOI was the body surface (mean=65%, Figure 4). The caudal fin contributed 18.4%, followed by the dorsal and anal fins (8.9 and 7.9%, respectively). The pelvic fin contributed very little to the total posterior MOI.

**Fig. 3.**
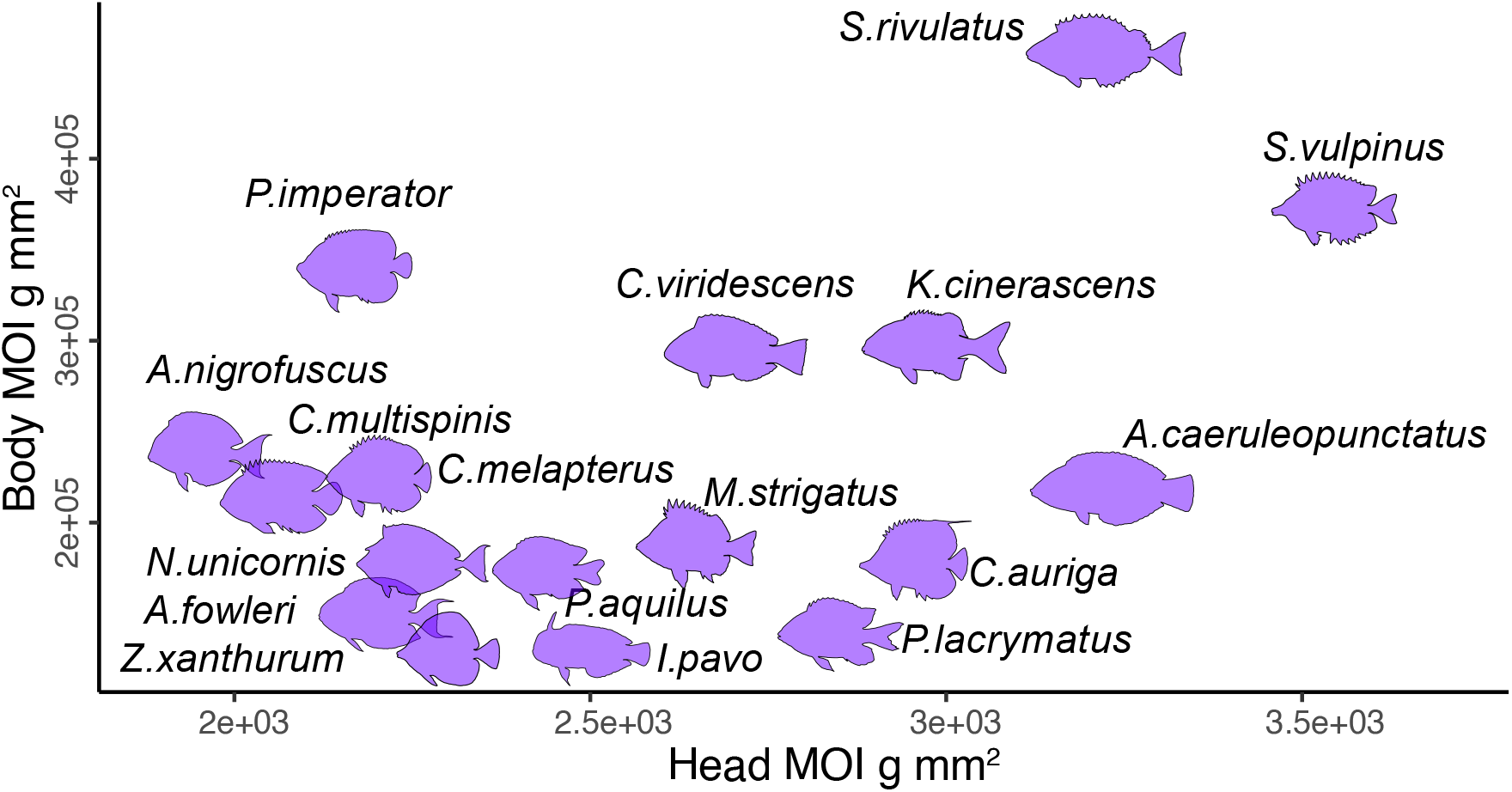
Morphological diversity. Models used in the experiment represented a range of morphologies, varying in the moment of inertia of the head, body, fins, and their combinations.

**Fig. 4.**
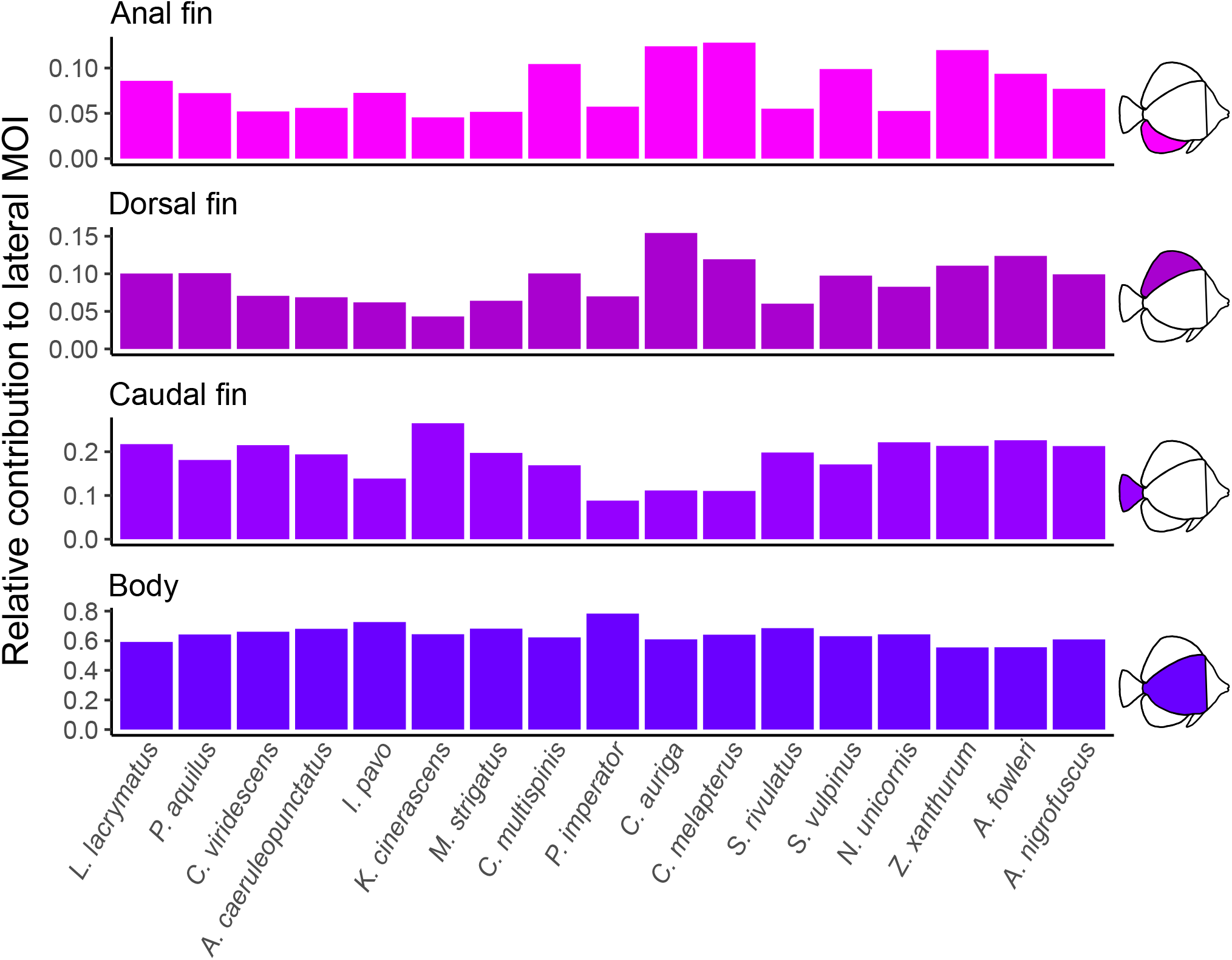
Relative contribution of lateral control surfaces. The total moment of inertia of the lateral profiles is a combination of the MOI of all lateral morphologies, including the body and the caudal, dorsal, and anal fins. In all models, the MOI of the body had the largest contribution to the lateral MOI while fin contribution varied between models. Note the different y-axis scale for each morphology.

The MOI of the caudal fin was correlated with that of the body (phylogenetically-corrected Pearson correlation coefficients = 0.686 and p-value*<*0.005), while the MOI of the anal fin was correlated to that of the dorsal fin (phylogenetically-corrected Pearson correlation coefficients = 0.915 and p-value*<*0.001). Head MOI was weakly correlated to the body MOI, but this correlation was not significant (phylogenetically-corrected Pearson correlation coefficients = 0.48 and p-value=0.06).

### Head flick speed is negatively correlated with head MOI

As hypothesized, the angular speed of the head flick was negatively correlated to the head MOI (Linear regression, slope = -0.25±0.03, p*<*0.001, whole model Adjusted *R*^2^ = 0.71, F(1,15)=39.9, p*<*0.001). The model with the lowest MOI had a mean angular speed of 1137 deg*/*s, while the model with the highest MOI had a mean angular speed of 799 deg*/*s (Figure 5A).

**Fig. 5.**
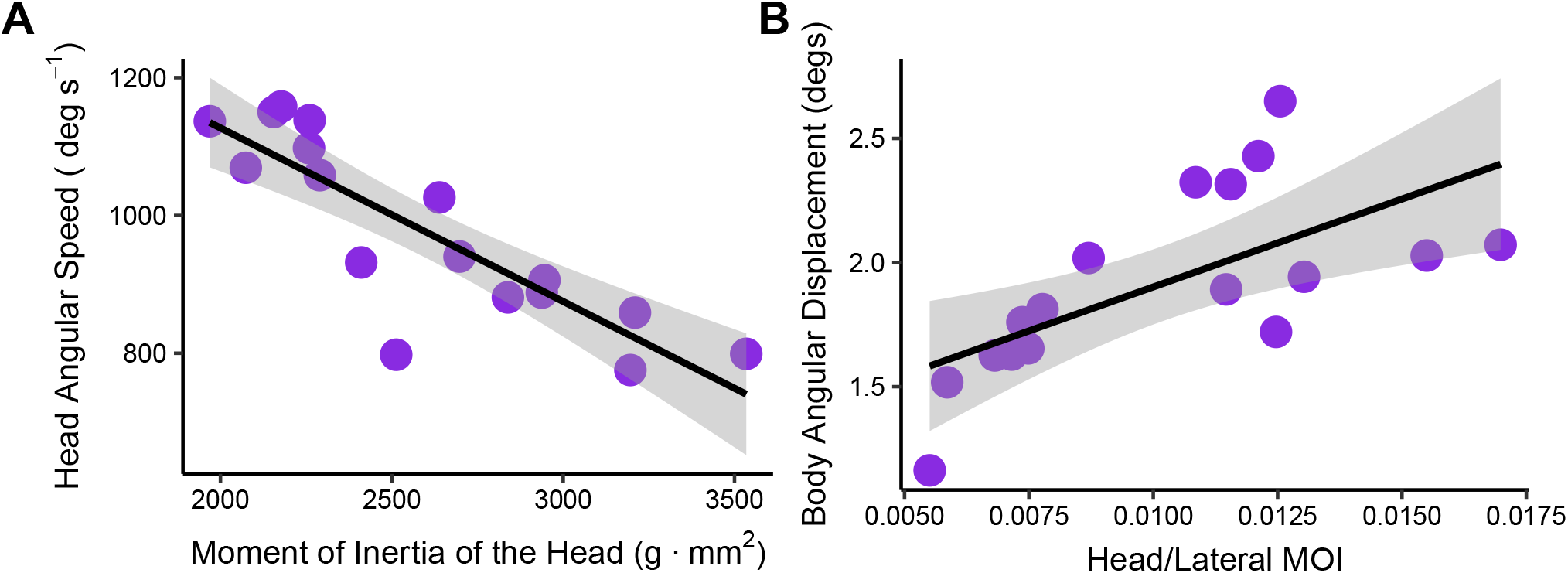
The effect of MOI on body and head kinematics. A) Heads characterized by lower MOI performed faster heads flicks (Linear regression, adjusted *R*^2^ = 0.71, p*<*0.001). B) Models with high MOI body and fins and low MOI heads displayed less angular displacement during head flicks (Linear regression, Adjusted *R*^2^ = 0.38, p*<*0.005). In both panels each point represents a different model (with y axis values denoting the average value across trials) and gray shading represents the 95% confidence interval.

### Body displacement during the flick is positively correlated with relative MOI

Body displacement during head flicks was positively correlated with the ratio between head and lateral MOI (Linear regression, slope = 70±21.44, p*<*0.005, whole model Adjusted *R*^2^ = 0.38, F(1,15)=10.92, p*<*0.005). The model with the lowest MOI ratio (0.005, smallest head MOI relative to the laterl MOI) had a mean displacement of 1.1 deg during the ∼0.04 seconds it took the model’s head to flick, while the largest angular displacement of 2.6 deg was observed in a model with an MOI ratio of 0.012 (Figure 5B).

## Discussion

Pulling from the substrate is a process that involves rapid rotational movements. Despite the prevalence of these rotational kinematics and their established contribution to feeding performance, whether and how head and body morphologies influence pulling kinematics has not been investigated. Here, we examined whether the moment of inertia of the head and body can be used as an informative index to describe and predict the relationship between morphology and kinematics. Using mechanical models representing the morphological diversity of pulling reef fish, we show that head morphologies that are characterized by low moment of inertia, and are less resistant to rotation, perform faster head flicks, thus facilitating pulling (Figure 5A). We further show that body stability during head flicks depends on the relationship between the morphologies of the head, body, and fins: Fish characterized with a high MOI body and median fins and low MOI head are more stable (i.e. display smaller displacement) during head flicks (Figure 5, B). These results provide the first mechanistic link between head and body morphologies, and the kinematics of feeding by pulling from the substrate.

### The usefulness of biomechanical models

The link between form and function is one of the most fundamental relationships in nature [Darwin, 1859]. Though it may seem as a straightforward relationship in many cases, the causal link between morphology, performance, and function may be challenging to demonstrate [Vogel, 1988, Wainwright and Reilly, 1994]. Biomechanical models are extremely useful for integrating multiple traits into a comparable quantitative framework. In such models, first principles are used to link morphology, often summarized through informative indices (e.g. lever ratio, drag coefficients, mass distribution), to the system’s kinematics, dynamics, or energetics. Well-fit models provide a quantitative relationship between morphology and performance, allowing an expansion of inference to unmeasured data based on the model’s estimates. Hence, an informative model exempts us from the tedious task of directly measuring performance and instead enables it to be estimated based on an established relationship. In suction-feeding fish, for example, the ‘suction index’ integrates muscle and buccal cavity measurements, which are easily obtained from specimens, into an informative index that is tightly correlated with suction performance [Wainwright et al., 2007, Holzman et al., 2008]. In labriform swimmers, the aspect ratio of the pectoral fin, integrating its length and area, is correlated not only with swimming performance but also predicts utilization of wave swept environments and the fish’s position in the water column [Fulton et al., 2001, 2005, Wainwright et al., 2002b]. Such indices, which allow us to predict both performance and ecology based on morphology alone, capture fundamental form-function relationships. The identification of such indices provides an important stepping stone in the study of organismal performance.

The moment of inertia of whole organisms, or specific moving morphologies, is a relevant index for understanding and predicting their kinematics. MOI has been examined in the context of flight kinematics in birds, bats, and insects [Rees, 1975, Thollesson and Norberg, 1991, Berg and Rayner, 1995], the spinning behaviors of cetaceans [Fish et al., 2024, 2006], and swimming kinematics of aquatic organisms [Dabiri et al., 2020, Porter et al., 2011], but has not yet been considered in the context of fish feeding and pulling from the substrate. This is in part due to the fact that the rotating kinematics of the head during pulling, and their influence on feeding performance, have only recently been directly measured [Perevolotsky et al., 2020]. Lateral movement of the mouth to pull substrate-attached prey requires the rotation of the whole head [due to the location of the joint between the skull and vertebral column, Perevolotsky et al., 2020]. This means that the entire morphology interacts with the water, influencing hydrodynamics and kinematics. Therefore, the moment of inertia, which incorporates the shape, size, and mass distribution, is a relevant index for predicting lateral movements through the water, such as those occurring during pulling. The same index used to identify morphologies that facilitate rotation can also be used for identifying morphologies that resist rotation, making it an informative index to evaluate the kinematic interplay between speed and stability occurring during pulling. Our results, linking the MOI of the head, body, and fin morphologies to pulling kinematics, highlight the functional role these morphologies play in the process of feeding from the substrate, suggesting a form-function relationship governing their evolution.

### Evolutionary implications

Body depth is one of the major axes of morphological diversification in teleost fish [Friedman et al., 2022, 2021, Martinez et al., 2021, Ghezelayagh et al., 2022]. Deep-bodied morphologies were shown to characterize demersal fish, reef fish, and biting fish, with herbivorous fish displaying elevated rates of body depth evolution [Larouche et al., 2020, Friedman et al., 2020, Borstein et al., 2019]. Similarly, rates of body shape evolution in biters were found to be almost twice as high as those of suction feeding fish [Corn et al., 2022]. This evolutionary signal hints at the selective pressures acting on these morphologies, suggesting a functional role yet to be identified. The most widely accepted hypothesis pertaining to function suggests that the increased projected lateral area, along with the narrow flexible body, facilitates unsteady swimming, turns, and maneuvers [Larouche et al., 2020, Webb, 1984a,b]. Along the same line, these morphologies were hypothesized to be advantageous for navigation through complex habitats, such as coral reefs, providing a potential explanation for the prevalence of these morphologies in reef-associated lineages [Langerhans and Reznick, 2010, Webb, 1984b, Di Santo and Goerig, 2025]. However, an empirical link between deep-bodied morphologies and maneuverability has been difficult to demonstrate [Webb, 1978, Schrank et al., 1999, Schrank and Webb, 1998, Webb et al., 1996]. For example, a recent study quantifying the variation in swimming behaviors of fish in the reef found no association between morphology and routine swimming behavior, implying that deep-bodied fish do not display more maneuverable swimming in their natural environment [Satterfield et al., 2023]. Our results suggest that deep-bodied morphologies, with large protruding dorsal and anal fins, increase the total lateral MOI (Figure 4) thus increasing resistance to undesired movements during feeding. The adaptive value of this newly quantified function might be reflected in the evolution of deep-bodied morphologies in coral reef fishes.

Deep-bodied morphologies are often coupled with large and prominent dorsal and anal fins, especially in reef fish. These large median fins are usually not used during routine swimming in reef-associated fish, which rely mostly on their pectoral fins for swimming and maneuvering [with the exception of tetraodontiform swimmers, Gerstner, 1999]. Protruding median fins, with large spines, play an important anti-predatory role. The increased lateral profile aids in exceeding the gape-limitation of potential predators, and, in teleost fish, spines of the dorsal and anal fins were shown to co-evolve with deep-bodied morphologies [Price et al., 2015]. In butterflyfish, solitary species with more exposed feeding strategies are also characterized by larger dorsal and anal spines [Hodge et al., 2018]. Here, we suggest an additional functional role for the protruding dorsal and anal fins, expanding the projected lateral area of the fish and stabilizing it during the rapid rotational movements characterizing pulling from the substrate. In fact, the contribution of the dorsal and anal fins might be underestimated by the MOI, because their mass is substantially lower than that of the body. Yet, despite the differences in mass and mechanical properties, the median fins act as a functional extension of the body. In addition, both the dorsal and anal fins usually extend all the way to the caudal peduncle and many times, especially in reef fish families, flare posteriorly. This posterior expansion of area, far from the center of rotation, has a substantial effect on the drag force acting on the rotating fish and slowing it. Notably, the thrust-based drag generated by dorsal and anal fins was considered to play a role in the accelerating movements of fish, especially in c-start maneuvering [Borazjani, 2013]. However, their role as stabilizers during feeding from the substrate has not yet been considered.

### A Potential Functional trade-off

Feeding performance, or the amount of food removed from the substrate during pulling, depends on the pull force exerted by the fish [Perevolotsky et al., 2020]. Increasing this force can be achieved either by increasing the rotational acceleration of the head or by increasing the torque applied on the prey. These solutions impose contrasting functional demands, and are facilitated by different morphologies [Dabiri et al., 2020]. Increasing torque can be achieved by extending the lever arm, or the distance between the mouth and the axis of rotation (i.e. head length, r). Increasing acceleration is achieved by minimizing the MOI of the head (therefore reducing r; Figure 5). These contrasting demands lead to a mechanical trade-off, where morphologies that increase torque (elongation of lever arm) also resist rotation and thus display lower acceleration. A potential morphological compromise might be that of a deep head at the base, which tapers towards the mouth. This shape reduces the mass far from the axis of rotation, thus reducing MOI, while still maximizing the lever arm length and increasing torque. Such head morphologies are actually quite common in reef fish, displayed by species from the Acanthuridae, Pomacanthidae, Siganidae, Chaetodontidae families, and more. To date, there is no consensus regarding the function of these tapering snout morphologies, with multiple hypotheses suggested, including an adaptation for crevice feeding and mating behavior [Fox and Bellwood, 2013, Brandl and Bellwood, 2013]. Our results suggest these tapering head morphologies might play a functional role in pulling from the substrate, facilitating both fast and forceful bites. Our experimental setup was designed to test the effect of morphology alone on head flick kinematics, but further work measuring or modeling the dynamics between speed and force during flicks might shed more light on this relationship.

### Limitations

Our experimental design allowed us to isolate the effect of morphology on kinematics, reducing the variance that originates from other variables. We show that the body acts as the largest passive stabilizing morphology and that overall, fish morphologies with higher MOI bodies relative to their heads stay more stable during head flicks. Notably, the displacement measured in this study was relatively small, especially when compared to observations of natural feeding events. This was a result of the short time frame at which kinematics was recorded (only during the head flick, less than 0.04 seconds long). Naturally, being propelled by the same spring but differing substantially in size and mass, the heads of models accelerated much faster than their bodies, reaching the 45 degrees of the flick while the bodies only moved a few degrees. We chose not to extend the kinematic analysis of body movement beyond the flick duration because the fully compressed spring would touch the body at the end of the flick, influencing its dynamics. Though the displacement is small, the clear trend that is captured demonstrates the functional advantage of specific body and head morphologies in the process of pulling from the substrate.

Our experimental design restricted the model’s movement to yawing, but it is possible that other undesired rotational movements (pitch and roll) would occur during head flicks. Rolling in particular is considered an undesired movement that fish constantly control by balancing buoyancy and mass [Webb, 2002, Di Santo and Goerig, 2025]. Fish can balance rolling actively by propelling the dorsal and anal fins or by contracting the muscles which protrude the fins, increasing their moment arm relative to the center of mass and center of buoyancy thus resisting rolling [Standen and Lauder, 2005, Webb, 2004]. Whether and how active fin expansion and kinematics play a role in stabilizing the body during pulling requires further investigation, although in-situ observations show expanded lateral fins during feeding [Figure 1, Perevolotsky et al., 2020, Mihalitsis and Wainwright, 2024]. Our mechanical fish were designed to isolated the effect of morphology on head flick kinematics, but other external and internal factors integrate to affect the fish’s motion underwater. Specifically, variation in muscle force and dynamics should have a dominant effect on the kinematics of the head and body, potentially overriding the effect of morphology. Furthermore, the variation in body girth may have a considerable effect on MOI, because body girth changes across the fish’s length. When applying our approach to real fish, the correct mass distribution should be used for calculating MOI. In addition, our model morphologies did not capture the inter-specific morphological variation, associated with age and sex [Herler et al., 2010, Echeverria, 1986]. Specifically, variation in fin morphology, a pronounced characteristic of sexual dimorphism, may lead to varying kinematic performance between males and females of each species.

Lastly, the intricate kinematics of the pectoral fins could not be replicated, and their role in stabilizing rotational feeding kinematics remains to be determined. That being said, and as described in [Perevolotsky et al., 2020], during head flicks the pectoral fins are engaged in propelling the body backwards. Thus, they cannot be actively utilized in stabilizing kinematics until this phase is over. This temporary restriction of the pectoral fins highlights the importance of other, passive, stabilizing control surfaces during head flicks. Though field and laboratory work with live fish under realistic flow regimes is needed, our results indicate that the lateral morphologies of both head, body, and fins of biting fish are functional morphologies that affect pulling performance. As such, they should be considered in studies of fish kinematics, ecomorphology, and evolution. We hypothesize that considering whole-body kinematics and morphology can shed light on large-scale selection-driven evolutionary processes shaping the ecomorphological diversity of biting fish.

## Acknowledgments

We thank Matthew Kolmann and the FHL fish class 2022: visitors, staff, and students, especially Duncan Kennedy and Finn Mander for technical and emotional support; Members of the Holzman lab and prof. Gal Ribak for insightful discussions regarding this project; The FHL and IUI communities, for creating a positive and supportive environment for science to happen.

## Author contributions statement

TP, JMB, CMD, APS, and RH conceived the project and methodology, TP performed the experiments, YR and YA digitized videos, TP and RH performed the analyses and wrote the initial manuscript. All authors contributed to the final manuscript

---

**Table 1.** Model Species

| Family | Species | Primary Diet | FishBase photo |
| --- | --- | --- | --- |
| Chaetodontidae | *Chaetodon auriga* | Facultative corallivore | Chaur_u7 |
| Chaetodontidae | *Chaetodon melapterus* | Corallivore | Chmel_ua |
| Pomacanthidae | *Centropyge multispinis* | Invertivore | Cemul_u7 |
| Pomacanthidae | *Pomacanthus imperator* | Invertivore | Poimp_u6 |
| Labridae | *Anampses caeruleopunctatus* | Invertivore | Ancae_u2 |
| Labridae | *Iniistius pavo* | Invertivore | Xypav_m1 |
| Labridae | *Calotomus viridescens* | Herbivore | Cavir_m3 |
| Acanthuridae | *Zebrasoma xanthurum* | Herbivore | zexan_u1 |
| Acanthuridae | *Acanthurus fowleri* | Herbivore | Acfow_u0 |
| Acanthuridae | *Acanthurus nigrofuscus* | Herbivore | Acnig_ua |
| Acanthuridae | *Naso unicornis* | Herbivore | Nauni_u1 |
| Siganidae | *Siganus vulpinus* | Herbivore | Sivul_u2 |
| Siganidae | *Siganus rivulatus* | Herbivore | Siriv_u3 |
| Pomacentridae | *Stegastes lacrymatus* | Herbivore | Pllac_u0 |
| Pomacentridae | *Pomacentrus aquilus* | Herbivore | Poaqu_u6 |
| Kyphosidae | *Kyphosus vaigiensis* | Herbivore | Kycin_u3 |
| Kyphosidae | *Microcanthus strigatus* | Herbivore | Mistr_u2 |

